# Architectural analyses of tree forking habit with single-image photogrammetry: a case study of post-mature temperate oaks

**DOI:** 10.1101/795286

**Authors:** Kamil Kędra

## Abstract

Tree forking is both ecologically and economically relevant, but remains much understudied. Here, thirty post-mature temperate oaks (*Quercus robur* or *Q. petraea*) forking habit was both qualitatively and quantitatively analysed with the single-image photogrammetry (SIP), in a north-exposed mixed, deciduous forest remnant (near Krakow; Poland). A new classification of mature oak architectures was proposed, based on the original Hallé-Oldeman model, with modified locations of the main branches and presence or absence of bifurcation in the main stem. Two of the new classes were most clearly represented by the studied oaks. It was found that the trees tended to either keep branches at varying heights, with no forks, or to iterate forking, with no major (non-fork) branches below the first fork. The quantitative analysis confirmed the applicability of the branch to parent stem diameter ratio to define a fork. Branching ratio was positively correlated with both tree diameter and height of a branch above the ground, which is consistent with a previous study, based on much younger trees. It is concluded, that most probably the tree-level factors and phenomena, such as water supplies and posture control, played the key role in the studied oaks forking habit. The SIP method enabled valuable insights into the large oaks’ forking, both at the tree and branch levels, and may be further employed to study mature trees’ bifurcation patterns. Based on this study, some possible improvements to the methodology were discussed.

## 1 Introduction

Bifurcation or forking is an important feature of tree branching systems, leading to the formation of two, more or less equivalent, axes instead of a single monopodial axis. Despite the large economical and ecological consequences, quantification of tree forking (TF) has gained surprisingly little attention from the research community (Colin et al., 2012). The cited paper seems to provide the first quantitative analysis of TF ever made, and in spite of rising both vital and intriguing scientific issues about TF, up to now it has not been discussed. The authors presented a study of young (up to 23 years old) sessile oak (*Quercus petraea* (Matt.) Liebl.) forking, in three oak plantations of varying initial stand densities. It was found that TF was most common in the lowest density site, and number of forks per tree increased with tree girth, tree height and age. The causes of forks (Barthélémy & Caraglio, 2007; Bell, 1991; Chaar & Colin, 1999; Collet et al., 2011; Hallé et al., 1978; Ningre & Colin, 2007), either shoot-level (traumatic) or tree-level (metamorphic), were not clearly distinguished. The authors concluded, that this was probably because the trees under study were too young to observe the final architectural patterns, and still, the traumatic TF causes dominated, or because those patterns were only weakly pronounced (Colin et al., 2012). Some other studies aimed at general classification of young trees as forked or not (Jensen & Löf, 2017; Kuehne et al., 2013), according to the predefined global tree models. However, both quantitative and qualitative analyses of TF are probably completely lacking in the case of large, old trees. This lag may be linked to the fact, that the rapidly developing remote sensing techniques for tree architectural inventory (Liang et al., 2019), has not yet been applied to analyse older trees’ bifurcation patterns.

Forking is most noticeable in tree species “normally” exhibiting a single, monopodial trunk. The distinction between sympodial or monopodial growth pattern is the key question in the famous Hallé-Oldeman (HO) architectural classification (Bell, 1991; Hallé & Oldeman, 1970; Hallé et al., 1978). Among the 23 HO models, elaborated to describe the diversity of tropical plants’ architectures (Hallé et al., 1978), the Rauh’s model (named after the German biologist Werner Rauh) seems to retain the largest monopodial tree species representation in Europe, encompassing taxa of such wide-spread genera as *Quercus, Pinus, Picea* and *Acer* (Fig. 1). Generally, the Rauh’s model describes light demanding and early-successional species. The trees of the complex *Quercus robur* L. / Q. *petraea* (Matt.) Liebl. (Gomory et al., 2001), here referred to as oaks (Q. *robur sensu lato*), are of major importance in forestry (Saenz-Romero et al., 2017); while being prone to forking (Colin et al., 2012), e.g., because of specific wood properties, such as the ability to form tortuous grain pattern, interlocking the forked junctions (Slater & Ennos, 2015). This and other traits, that contribute to tree plasticity in relation to local growth environment, make the original architectural “blueprint” (Rauh’s model) hardly recognisable at the scale of whole mature and older oak trees (Oldeman, 1990). Therefore, it is worthwhile to test whether any other architectural pattern or patterns may be useful for describing post-mature oak trees.

**Fig. 1.**
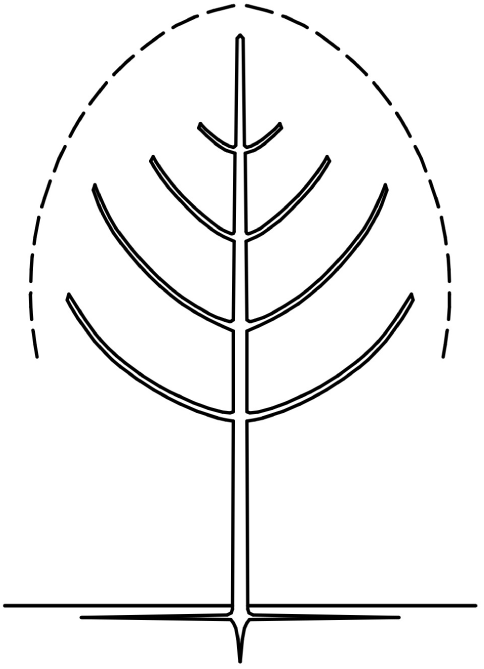
A simplified Rauh’s model (Hallé et al., 1978), showing its main characteristics: monopodial trunk and rhythmic, orthotropic branching

This study was conducted in an old-growth, oak-lime-hornbeam, north-exposed small forest remnant, where many of the target oak trees exhibited forking, to provide both tree- and branch-level assessment of TF habit. The following objectives were addressed:

1. Define and test the possible architectural patterns of mature and old oaks, including forking habit, as a modification of the original (Rauh’s) model;
2. Develop new ways to quantitatively analyse and describe large trees’ forking, with an image-based remote sensing method.

## 2 Methods

### 2.1 Study area and tree sampling

The site (Krzyszkowice Forest) is located within a local hill (ca. 120 ha area, ca. 65 m relative height: between 220 and 285 m a.s.l.), in the vicinity (SE) of Krakow, Southern Poland (50°0′3.14″N 20°0′41.2″E). The loess-mantled hill has an elongated shape, along the longitudinal direction; with the northern (larger) and western (smaller) slopes covered with the forest (ca. 34 ha), while the southern and eastern slopes are mainly covered with discontinuous urban fabric and agricultural areas (Urban Atlas 2012: <https://land.copernicus.eu/local/urban-atlas/urban-atlas-2012>). This is one of the few old-forests within the urbanized district; those forest remnants mostly occur in the form of small, isolated “islands”, and usually subjected to some kind of nature or landscape protection. The Krzyszkowice Forest is under a partial protection (since 1998) to preserve the mixed, deciduous oak-lime-hornbeam forest (*Tilio-Carpinetum*), and valuable fauna and flora, including relict mountain plants’ locations (Gazda & Gazda, 2010). The loess-mantled soils are fertile, and prone to erosion, as indicated by two distinct gullies within the forested area. This site was previously described, in less detail, in the conference paper by Kędra et al. (2016), focusing on inter-tree competition, and other external factors influencing oaks’ crown radii and the overall crown size and shape.

The whole site is covered with a network of permanent, circular plots (0.05 ha each), regularly spaced (see Kędra et al. (2016) for a map). The plots were established in 2007 (Gazda Anna, personal communication), and are individual tree-centred, with 30 plots targeting at mature oaks, with diameter at breast height (DBH) larger than 40 cm. Those trees were used in this study, as well as in Kędra et al. (2016). Here, however, the oaks were classified as being of the *Quercus robur/Q. petraea* complex, instead of *Q. robur sensu stricto*. This more general approach is correct in the presence of both oak species within the forest, their frequent hybridization (Gomory et al., 2001), and lack of genetic identification of the individuals. The target trees median DBH was 53 cm. The exact age of those trees is not known; however, they are estimated to well exceed one hundred years in age. The neighbourhood (other trees and shrubs growing within the plots) included several deciduous tree species, mainly: silver birch (Betula pendula Roth), common hornbeam (*Carpinus betulus* L.), oaks, sycamore maple (*Acer pseudoplatanus* L.), small-leaved lime (*Tilia cordata* Mill.), and wych elm (*Ulmus glabra* Huds.). The mean density of trees and shrubs (with DBH of at least 7 cm) was 24±7 individuals per plot (480±140, up-scaled to individuals per hectare), and mean basal area was 1.75±0.46 m^2^ per plot (35±9 m^2^/ha). The plots were characterized by varying local terrain slopes (3.5±3.3°).

### 2.2 Image acquisition

The single-image photogrammetry (Gazda & Kędra, 2017; Kędra et al., 2019), requires one, high resolution photograph per tree. At the moment of image taking, the whole branching system is flattened at once, in relation to a theoretic “projection plane”. Therefore, the place from which the image is to be taken, needs to be carefully selected. The main concept is to capture the representative crown profile; this usually means to capture the largest crown asymmetry, or the largest crown horizontal extent (if the tree of interest was not visibly inclined in any direction). Here, the following protocol was utilized: first, examining the tree crown from below (standing next to the stem) to determine the major direction of the crown development. The trees were never perfectly symmetrical and it was always possible to point such direction, and note the azimuth. Second, subtracting and adding 90 degrees from and to the azimuth (respectively) to determine two possible directions of the image to be taken. Third, choosing between the two possible (opposite) image directions, to assure the best possible visibility (lowest occlusion) of the whole branching system. Finally, taking the photograph, with specific settings: distance from the target tree and camera tilt (keeping in mind that decreasing the distance and increasing the tilt angle may negatively affect the measurement accuracy (Gazda & Kędra, 2017)). The noted settings were further used to transform the images from non-metric to metric ones, with the QGIS software v.2.8.9 (QGIS Development Team, 2016).

### 2.3 Qualitative analysis

The original Rauh’s model (Fig. 2, R.A) implies that the main tree axis (monopodial trunk) contributes to the vertical tree extent, while the lowest branches (of equal insertion height) are the main branches, that contribute to the horizontal crown extent. Three variants of the original model were developed (Fig. 2, R.B-D), which account for modifications in the main trunk (forking) and/or the main branches’ insertion heights. The model R.B maintains the monopodial trunk, but the main branches are of unequal insertion height (e.g. because of branch mortality in low light conditions). The models R.C and R.D both account for TF, however, in the former, only one of the two main branches is affected by forking (the other branch is located below the fork); while in the latter, both main branches (and the whole crown) are affected by TF. The architectures of the target oaks were carefully analysed, with the use of photographs taken (as described in the previous section), and according to the presented models (R.A-D).

**Fig. 2.**
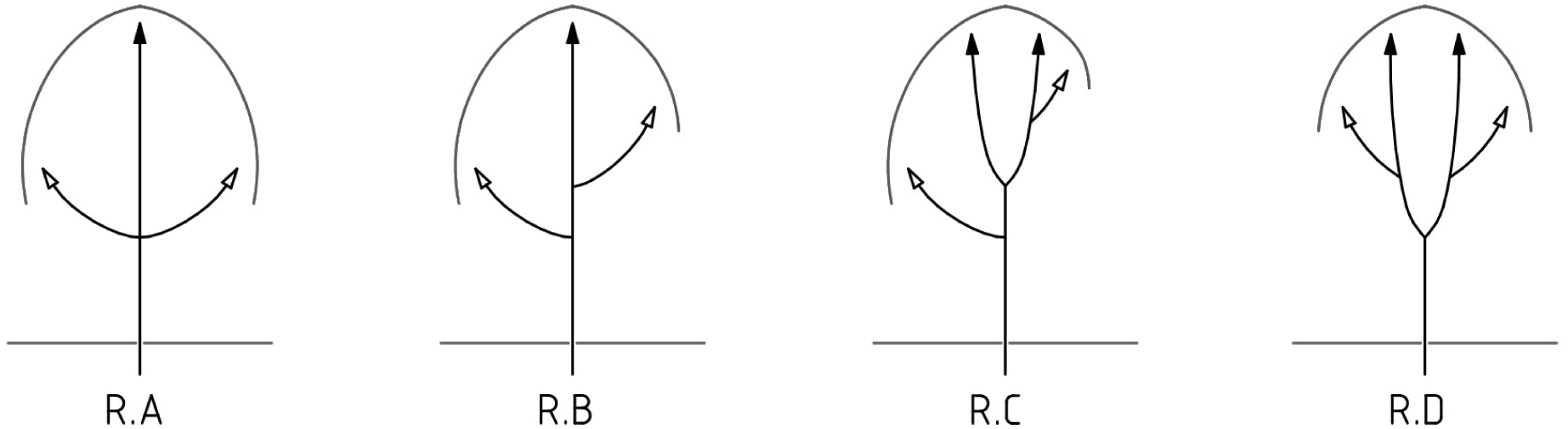
The original Rauh’s model (R.A), and its modifications (R.B-D), black arrows denote axes that contribute to the vertical tree extent, and the empty arrows denote main branches, which contribute to the horizontal crown extent

### 2.4 Quantitative analysis

#### 2.4.1 Extraction of the candidate fork-related variables

##### Branch sampling

Thirty trees were under study; therefore, the aim was to analyse 60 main branches (two per tree), in terms of the variables (architectural traits) that may potentially describe and define branches as forked or not in a quantitative manner. Only the main branches were considered, that is, those which contributed to the horizontal crown extent, within the selected projection plane. The set of considered traits included: (1) branch diameter (or branch thickness, BT), (2) the corresponding main stem diameter (or stem thickness, ST), (3) the ratio between BT and ST (branching ratio, BR), (4) branch insertion point height above the ground level (branch height, BH), branch angles: (5) in relation to the main stem position (relativeBA), (6) in relation to the vertical direction (absoluteBA), (7) the difference between relativeBA and absoluteBA (deltaBA1), and (8) the modulus of deltaBA1 (deltaBA2). The measurements of the traits (1-4) were taken with the ArchiCAD software <https://myarchicad.com/>, and the branching angles were measured in the LibreCAD (open source) software <https://librecad.org/>.

##### Branch and stem diameters

The diameter measurements, including DBH, were analogous to Kędra et al. (2019); however, here the branch and parent stem diameters were measured at the distance of 1 m from the branch insertion point, instead of the 0.5 m distance used in that study (because the trees here were larger, with thicker branches: the DBH of target trees here was approximately twice larger than the DBH of that study’s trees).

##### Branching ratio

The relation between the branch diameter and the parent stem diameter (branching ratio, BR) has been used to quantitatively define a fork (Colin et al., 2012; Ningre, 1997). BR may take a range of values: between more than zero and one; BR lower than 1/2 denotes a small branch; BR between 1/2 and 2/3 denotes a large branch; and BR larger than 2/3 defines a fork, while the values closer to the 2/3 threshold reveal asymmetric forks, and BRs close to 1:1 indicate a true fork, resulting in two equal axes (Ningre, 1997). Here, branching ratios were calculated for all branches that contributed to horizontal crown extent and underwent diameter measurements.

##### Branch height

Colin et al. (2012) found that higher trees may have more forked junctions in the branching system, than the lower trees; therefore, it was suspected that branches located higher in the tree were more prone to forking than the lower-located branches. Herein, the branch height (BH) was measured as the vertical distance between each tree’s base point and each branch axis intersection point with the main stem axis (branch insertion point). It is stressed, however, that BH cannot be regarded as the height of the first (lowest) branch, which has been used in several studies as the location of live-crown base, e.g. (Burkardt et al., 2019), because some minor (but vital) branches could be present, below the main branches, in case of each analysed tree.

##### Branch angles

Generally, forked branches are thought to have a more upright position than the non-forked branches. Branching angle measurements have gained much attention from the research community, dealing with remote sensing of tree architecture (Bayer et al., 2013; Burkardt et al., 2019; Kędra et al., 2019; Pyorala et al., 2018). This type of traits proved to be useful for examining the effects of tree species mixing (Bayer et al., 2013). However, it seems that there is no widely accepted protocol on branch angle mensuration. Several ways to measure this trait were proposed, including the relative angle, between the branch and the parent stem (Kędra et al., 2019), and the “absolute” angle, between the branch direction and the vertical direction (Bayer et al., 2013). Here, both the relative and absolute branch angle measures were used, as well as the differences between both of them (which represent the level of local inclination of the main stem), to see which of those traits has the highest potential to discriminate a forked from a non-forked branch.

#### 2.4.2 Statistical methods

The variables were examined according to the standard methods for probability distribution estimation (histograms and probability density curves). The traits were split into two groups, with regards to forked or non-forked branches, as defined in terms of the qualitative assessment. To test whether those traits may differentiate the forked branches from the non-forked ones, a Kruskal-Wallis nonparametric test was used, with the null hypothesis stating that both groups of each variable come from populations with the same distribution. This test requires homoscedasticity in the data, and this was checked with the Levene’s test, in the R package “car” (Fox & Weisberg, 2019). When heteroscedasticity was found, Welch’s test for heteoscedastic data was used. Finally, the correlation analysis was performed to determine the monotonic relationships among the architectural variables and between those traits and the general measure of tree size, here represented by DBH. The results were plotted with the use of the “corrplot” package in R (Wei & Simko, 2017). All statistical analyses were performed with R v.3.2.3 or v.3.4.1 (R Core Team, 2017).

## 3 Results and discussion

### 3.1 Qualitative results

The main criterion of the proposed architectural classification was whether the analysed main branches were fork-related or non-fork-related. Keeping only this is mind, almost all the trees could be definitely assigned to the presented models: R.A-D (except for a single tree, see Fig. S1, model R.C, tree number 5). However, when considering the second criterion in the models R.A and R.B, i.e. constantly monopodial stem above the two main branches, some trees failed to be assigned to those models, as there were considerable bifurcations in the upper part of the stems. Therefore, two “forked” submodels were added: R.Af and R.Bf, to include those trees in the general classification. Not a single tree fully conformed to the original Rauh’s model, and only two trees had the two main branches with overlapping bases, while exhibiting forking of the main axis above the main branches (Fig. 3). The three models: R.B, R.C and R.D, were similarly represented (between 23 and 30% of all trees). However, four other trees were classified under the R.Bf model, and when those trees were pooled together with the R.B model trees, then this architectural type (fork-unrelated main branches, at varying heights) dominated considerably (40% of all trees). Five trees (55%) of the R.D model had some minor or dead branches (seemingly once being major branches) below the fork (Fig. 4a), suggesting that the R.C model trees might present a tentative state (which may change to R.D model in the future). Furthermore, the R.Af and R.B/Bf trees could also turn to the R.D model with time, as exemplified by one distinct reduction of the main tree axis in the presence of two main branches close to each other, leading to the formation of a pseudo-fork (Fig. 4b). On the other hand, forks may also be reduced to single axes, which was observed in the case of a single tree, low in the crown (6.2 m above the ground, which was the minimal BH of this study; Fig. 4c). Many of the R.D trees exhibited repeated forking of the axes coming from a fork below. This may suggest that there were some global, tree-level factors, underlying this forking habit, rather than single shoot-level traumatisms. Supplementary Figure 1 presents all trees, and their classification to the qualitative models (summarized in Tab. 1).

**Table 1.** Summary of the qualitative results and branch sampling

| Model | Number of trees | Number of branches | Measured branches | % measured branches |
| --- | --- | --- | --- | --- |
| R.Af | 2 | 4 | 0 | 0 |
| R.B | 8 | 16 | 12 | 75 |
| R.Bf | 4 | 8 | 6 | 75 |
| R.C | 7 | 13 | 11 | 85 |
| R.D | 9 | 12 | 8 | 66 |
| Total | 30 | 53 | 37 | 70 |

**Fig. 3.**
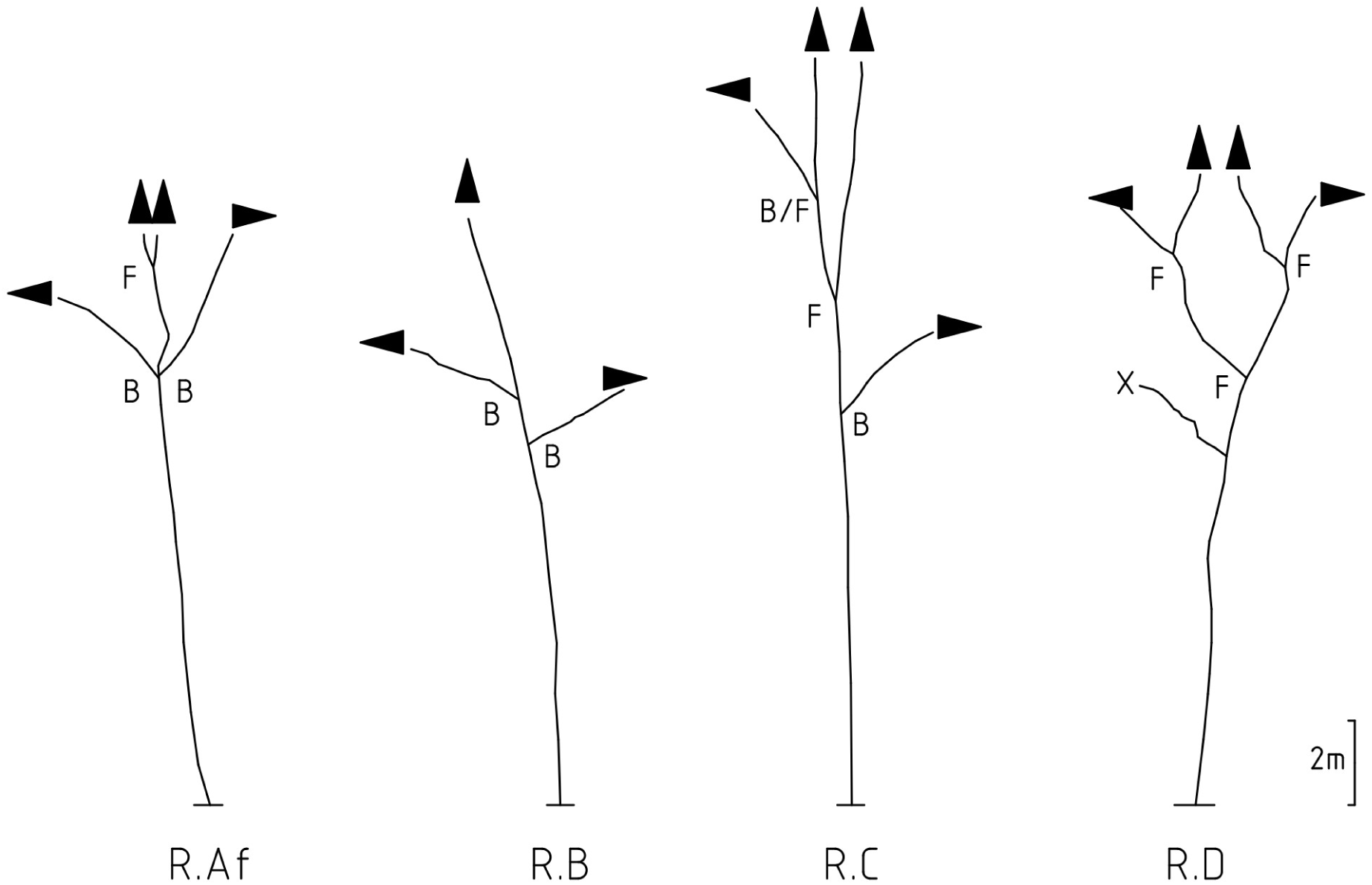
The most typical examples of actual branching systems, digitized with SIP, conforming to the proposed qualitative models (R.A-D); only the main axes and main branches were shown; B=non-forked branch, F=forked branch, X=dead branch

**Fig. 4.**
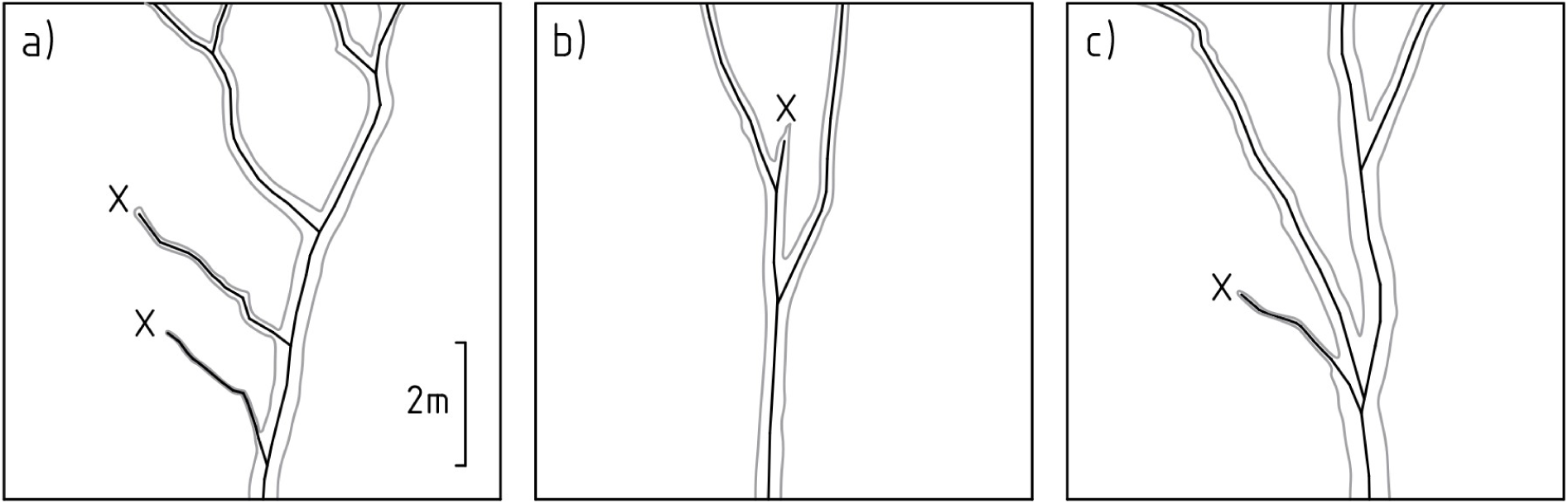
Three different types of axes reduction (X): a) reduction of branches below a fork; b) pseudo-fork formation by reduction of the main axis; c) one of the axes that once formed a fork was reduced, instead, a branch was formed later (next to the right); the scale is constant for a)-c)

### 3.2 Quantitative results

Fifty three branches were analysed in detail (see Fig. S2); seven cases were excluded from this analysis, because there was no considerable, or well visible, ramification of the main axis; in such cases it was assumed, that the main axis contributed both to the vertical and horizontal crown extent. Further 16 branches were excluded from measurements, mostly because there was another branch (or branches) within the distance of 1 m from the target branch insertion point, or the analysed branch was occluded by another one or the stem. In several cases the measurement was preceded by adjustment of the measurement radius (by 10 or 20 cm). Finally, a set of 37 branches underwent measurement of all described traits (Tab. 1); which included 23 non-forked branches and 14 forked branches (as determined qualitatively).

Most of the variables had notably right-skewed distributions (Fig. 5). The most bimodal-like distribution was found in the case of branching ratio (BR, Fig.5c); this might suggest that the mechanisms underlying forked and non-forked branches formation differ. Obviously, BR was the best fork/non-fork-disentangling trait; confirming that BR is a suitable fork-defining variable. Furthermore, slight symptoms of a second peak in the probability distribution curves were observed in case of branch diameter (BT, Fig.5a), relative branch angle (relativeBA, Fig.5e) and deltaBAs (Fig.5g,h).

**Fig. 5.**
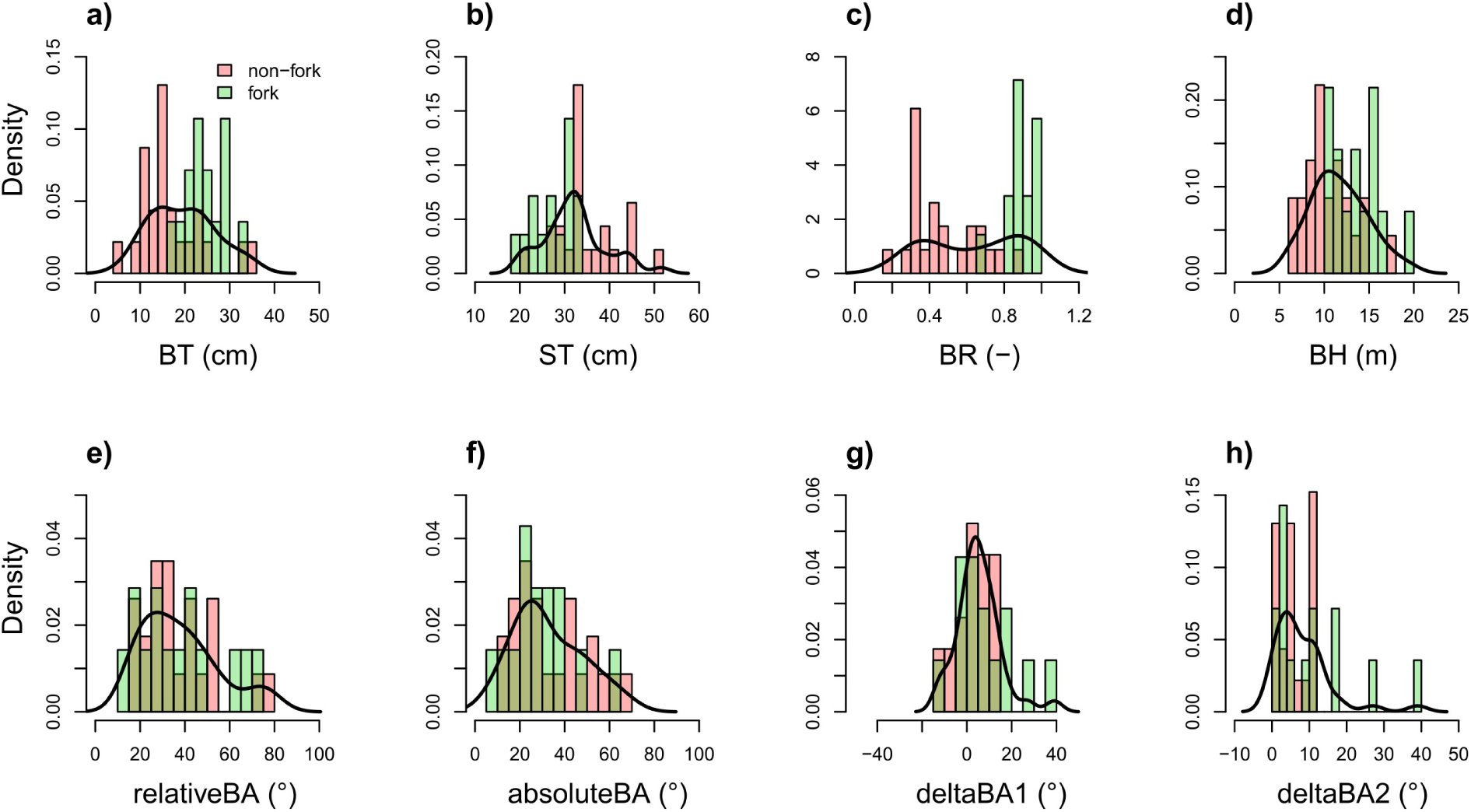
Histograms and probability density curves for the analysed architectural traits; bins corresponding to the forked branches are in green, and the bins for non-forked branches are in red; the fills are partly transparent to show whether they overlap or not

The Levene’s test revealed heteroscedasticity in case of BR and deltaBA2, while deltaBA1 was close to violation of the Kruskal-Wallis test’s homoscedasticity assumption (Tab. 2). Therefore, Welch’s anova test was used to look for differences between fork and non-fork groups of that variables. This was found in case of four traits, namely: branch diameter, stem diameter, branching ratio and branch insertion height (Fig. 6a-d). Interestingly, none of the angular measures significantly differentiated the forked from the non-forked branches. This was probably because of the relatively short distance, at which the angles were measured (1 m, in most cases), when the forked branches reached their upright positions further away from the branch insertion points.

**Table 2.**
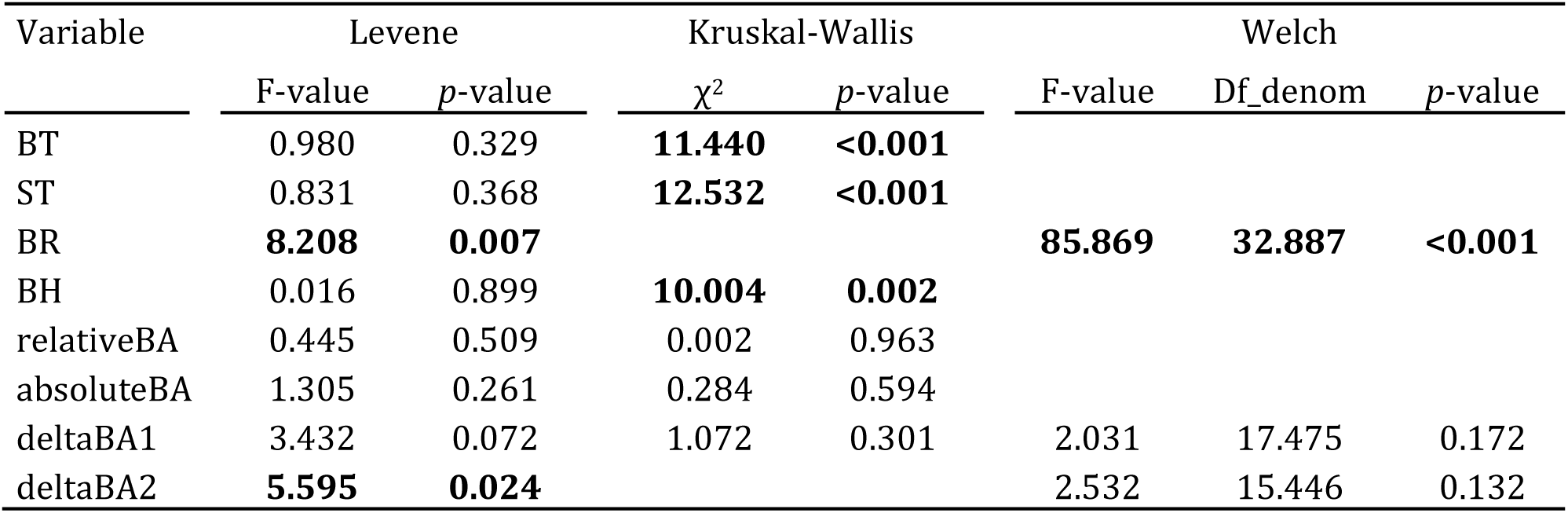
Results of the Levene’s, Kruskal-Wallis and Welch’s tests; Df_denom is the degrees of freedom denominator; the statistically significant differences were denoted by bold font

| Variable | Levene |  | Kruskal-Wallis |  | Welch |  |  |
| --- | --- | --- | --- | --- | --- | --- | --- |
| | F-value | p-value | $\chi^2$ | p-value | F-value | Df_denom | p-value |
| BT | 0.980 | 0.329 | **11.440** | **<0.001** |  |  |  |
| ST | 0.831 | 0.368 | **12.532** | **<0.001** |  |  |  |
| BR | **8.208** | **0.007** |  |  | **85.869** | **32.887** | **<0.001** |
| BH | 0.016 | 0.899 | **10.004** | **0.002** |  |  |  |
| relativeBA | 0.445 | 0.509 | 0.002 | 0.963 |  |  |  |
| absoluteBA | 1.305 | 0.261 | 0.284 | 0.594 |  |  |  |
| deltaBA1 | 3.432 | 0.072 | 1.072 | 0.301 | 2.031 | 17.475 | 0.172 |
| deltaBA2 | **5.595** | **0.024** |  |  | 2.532 | 15.446 | 0.132 |

**Fig. 6.**
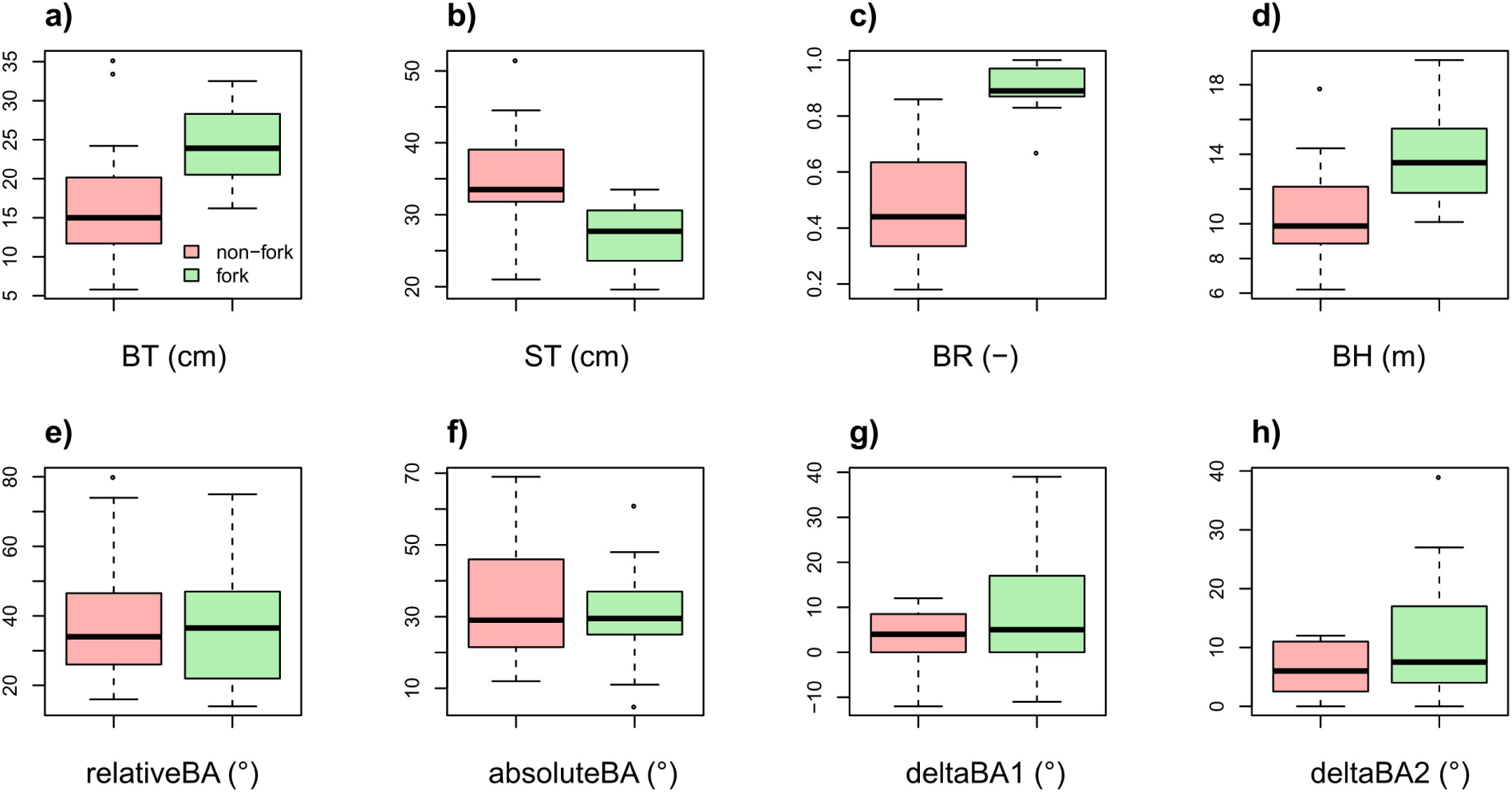
Box plots for the analysed architectural traits; boxes corresponding to the forked branches are in green, and the boxes for non-forked branches are in red; the plots represent the following statistics: minimum, first quartile, median, third quartile, and maximum

The Spearman’s correlation analysis (Fig. 7) showed that the branching ratio was positively correlated with tree size (DBH; rs=0.378, *p*=0.021), branch diameter (BT; rs=0.833, *p*<0.001) and branch height (BH; rs=0.587, *p*<0.001); while it was negatively correlated with the parent stem diameter of a branch (ST; rs=-0.576, *p*<0.001). The fact that fork prevalence, in the studied oaks, increased both with tree size and height of branches above the ground level, remains in agreement with the previous study of Colin et al. (2012), based on much younger trees. Furthermore, an insight into why branching ratio well describes forked branches was provided: the forked branches were thicker than the non-forked ones, while the corresponding main stem was generally thinner in case of the former branches (which may be linked to the observation, that they occurred higher in the stem); after all, the ratio between BT and ST only emphasized the differentiations provided by the both variables alone. The angular measures were not significantly correlated with BR; however, it is noticed that absoluteBA and deltaBA1 were seemingly more related with BR than relativeBA and deltaBA2. The former mentioned traits may probably become more useful in such analysis, with the way of measurement modified as described above.

**Fig. 7.**
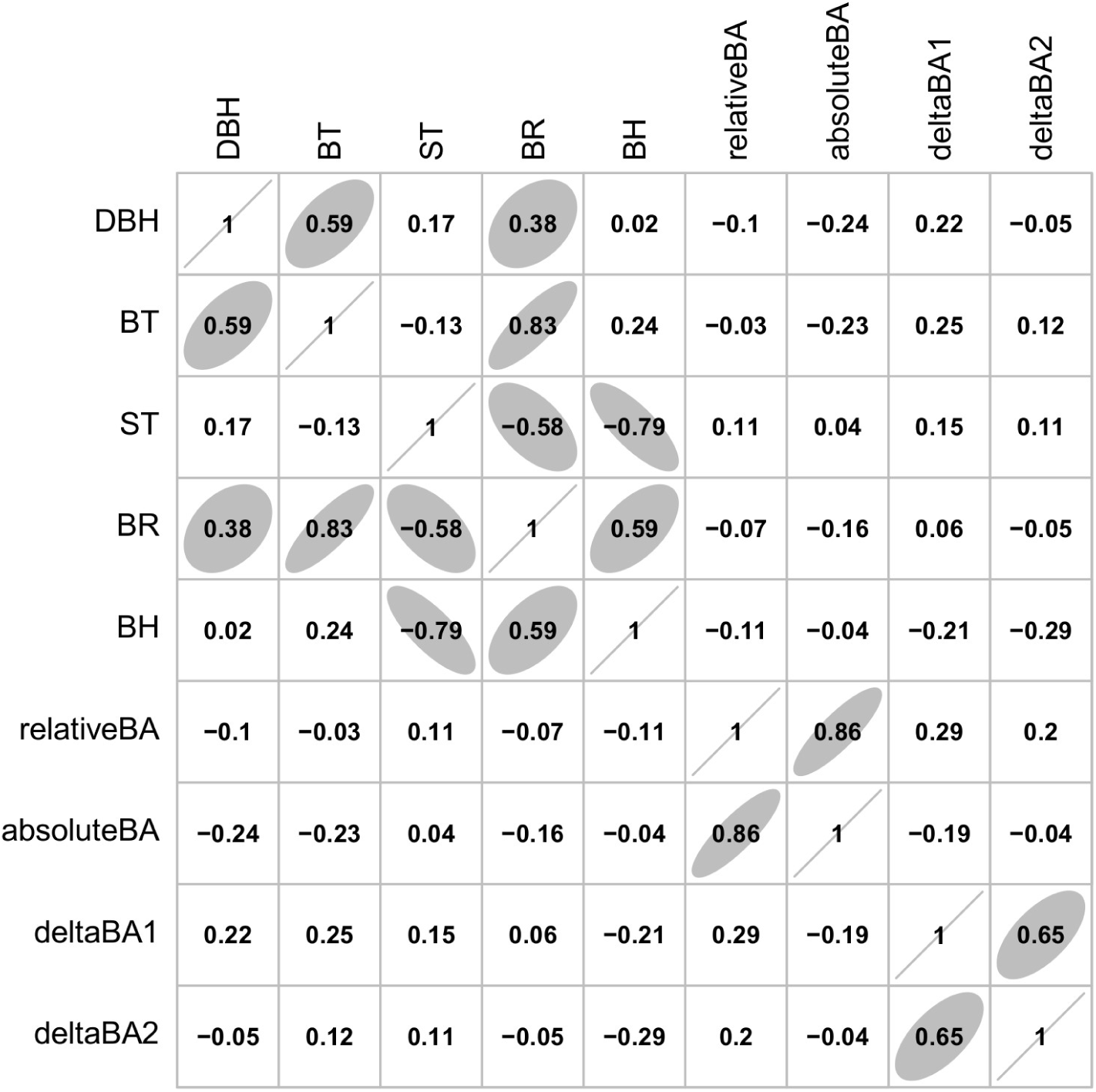
Results of the Spearman’s correlation analysis; statistically significant relationships (at the 0.05 level) were visualized with ellipses

### 3.3 Comparison of the proposed architectural classification with previous studies

Recently, several classifications were proposed to identify some general architectural patterns in young oaks (Jensen & Löf, 2017; Kuehne et al., 2013). Those models were developed strictly to assess the wood quality of the future harvest trees. Tree forking was explicitly included in the classification by Kuehne et al. (2013), which consisted of four classes: monopodial, steeply-angled, forked and brushy trees. The classification by Jensen & Löf (2017) included forking only implicitly, in the dipodial class (the other two possible classes were: monopodial and multipodial). Nonetheles, the two classifications seem quite similar to each other, they also both account for the level of curvature in the main stem. In comparison with that studies, the classification proposed here is more complex, because it requires selection of particular branches or axes, that contribute to the horizontal crown extent in relation to a certain vertical plane (which also needs to be defined). However, the calssifications by Kuehne et al. (2013) and by Jensen & Löf (2017) principally focus on the main stem (i.e. whether branching affects the stem or not), while in the classification presented herein, the main focus is on the branches: their relative position and whether they are affected by forking or not. Therefore, the selection between the methods mentioned above would depend on the purpose of any possible study: if tree forking is to be assessed in more detail, then the proposed classification seems appriopriate (with four different types of forked trees, namely models R.Af, R.Bf, R.C and R.D).

### 3.4 Qualitative vs. quantitative fork detection

In the qualitative TF analysis, when no measurements were taken, the shape of considered axes was crucial for the classification of a single ramification as forked or not. Most commonly, a forked branch was accompanied by a distinct, curvilinear shape of the other axis (Fig. 8, left). This pattern undoubtedly represented a fork (when both axes were vital, see Fig. 4c for an opposite example). However, for five branches, in trees of the R.B/Bf or R.C models (see Fig. S2), the quantitative analysis revealed that despite the main axis was clearly vertical (Fig. 8, right), the branching ratio well exceeded the 2/3 threshold (for two such branches it was higher than 0.8). It is known, that a branch may “escape” apical control in monopodial species (Groover, 2016); and oaks, because of the high wood density and firm branch attachment, are able to maintain very thick branches, growing horizontally from the vertical stem. Here, I propose not to classify such branches as forks, because they have no (or little) impact on the main axis shape. This implies that the qualitative analysis was more robust than the quantitative one. Nonetheless, agreement between both methods was rather high (87%), and branching ratio may still be recognised a simple and useful measure of TF. Furthermore, inclusion of the branch and stem shape metrics (Moulia et al., 2019; Moulia & Fournier, 2009) in the quantitative analysis could potentially resolve described ambiguities.

**Fig. 8.**
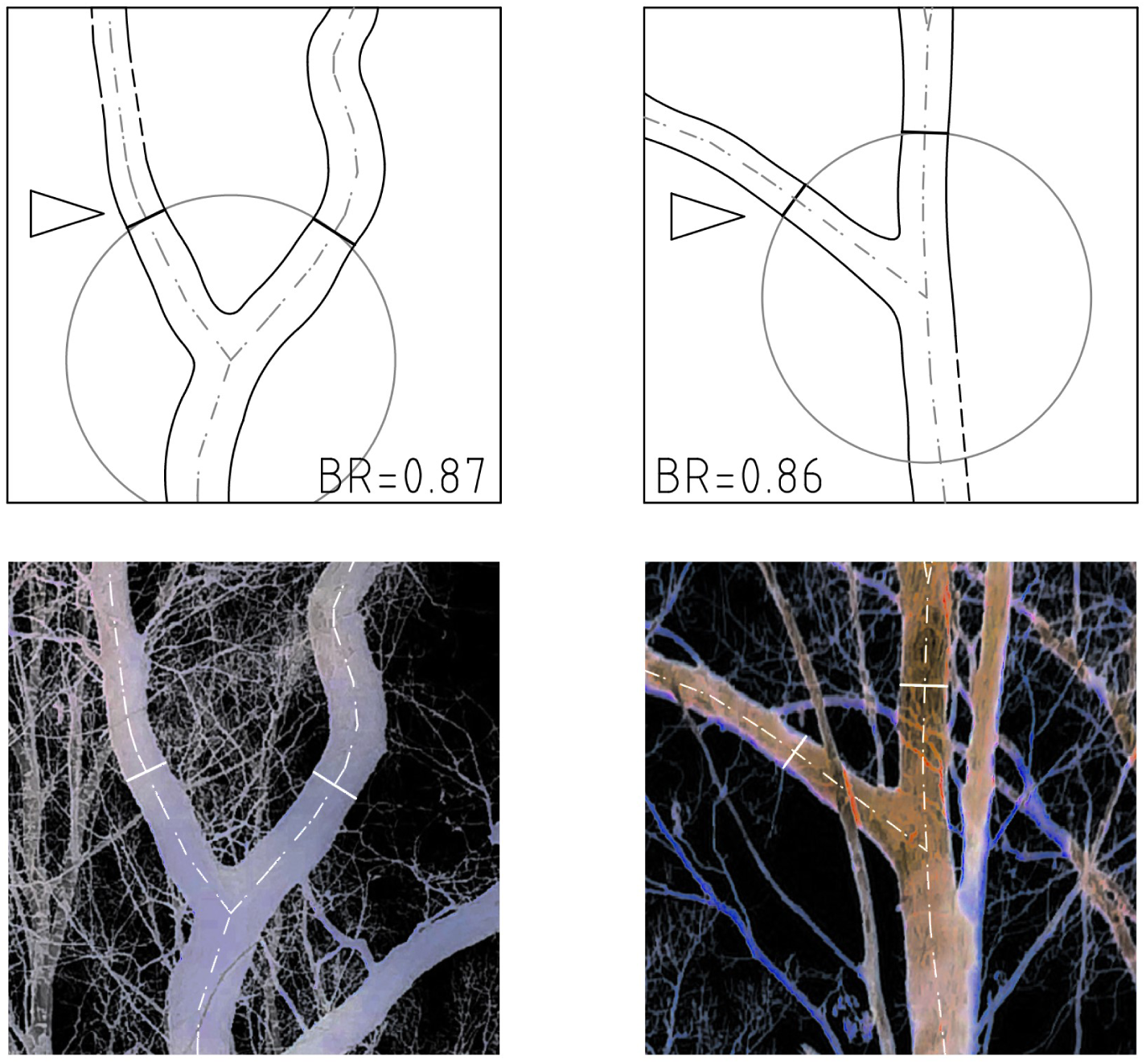
Two oaks’ branching outlines (top) extracted from images (bottom; with inverted luminosity); left: fork-qualified, right: non-fork qualified; quantitatively, in terms of the branching ratio (BR), the two junctions are roughly the same

### 3.5 Remarks on fork formation

Definitely, a separate, comprehensive review paper on fork formation in trees is needed; probably written by an interdisciplinary group of scientists dealing with plant growth and development, biomechanics, hydraulics and architecture. Herein, only some preliminary remarks were made, in relation to the presented results.

Several cues point to a general finding, that the tree-level causes of fork formation dominated here over the shoot-level (traumatic) ones. First, different factors and phenomena, that act globally on a tree, may result in a “forked” architecture. In the Leeuwenberg’s HO model, the equivalent branching is influenced by the terminal flowering (Hallé et al., 1978). In the shade-adapted architectural model (Pickett & Kempf, 1980), a similar pattern is formed because of low light conditions (and therefore, wide branching angles) within a forest understory. However, oak trees exhibit lateral flowering (not affecting branching). Moreover, the general light conditions here were rather homogeneously low, within the northern slope of the hill (although, the particular past growth conditions are not known). It is suspected that the variations in local topography, and thus, ground water availability (Radula et al., 2018) contributed here to the variation in the observed architectural patterns. Changes in water status may largely impact branch and stem diameter growth (Dietrich et al., 2018). Here, the measured diameters best disentangled the investigated branching habits. Second, trees control their posture as a whole (Moulia et al., 2019), and as they grow large, with heavy branches, the posture control must be an important issue. Stable patterns in branching systems may be found in even much older trees than those studied here (Dassot et al., 2019). Third, the apical control (a mechanism that promotes a single leader shoot) may be weakened in older trees (Wilson, 2000). Finally, the traumatic fork origins, such as frost damage (Ningre & Colin, 2007) or herbivory, mainly affect young (and short) trees, staying close to the ground. In this study’s canopy trees, only one tree displayed a clear (relatively recent) traumatism of the main axis, which led to forking (Fig. 4b), probably after a strong wind event. It seems that in most cases here, the forked junctions had been formed since the early times of branch emergence, as indicated by the distinct, curvilinear shape of both axes constituting the fork (Fig. 8, left).

### 3.6 Considerations for future studies

Based on this study, several aspects of the presented methods could be modified in future investigations, not to reproduce some weaknesses revealed here, while other aspects deserve endorsement. Firstly, it is clear that in such old and somewhat crooked trees there is little chance to measure both main branches in case of every tree, following the presented methods. Furthermore, even if most of the target trees underwent full measurements, there would still be a rather problematic nesting in the obtained data. Every two branches belonging to a particular tree are not fully independent from each other, and a group of two observations is much too small to be accounted for, e.g. in terms of random effects, by any modelling procedure. Therefore, it is suggested to either solely focus on a single main branch, or additionally select at least four other branches to be measured, in case of every tree. Those additional branches should be placed approximately within the same vertical plane as the main branch. To facilitate the workflow and to decrease uncertainty, it seems useful to mark all selected branches on the image taken, directly in the field (e.g. after opening the image with a portable tablet device). Secondly, the branch shape metrics could be improved to better represent crooked branches; this might be achieved by increasing the number of angle measurements per each branch (i.e. measuring branch angles at several distances from the branch insertion point). Thirdly, it is noted that the (free and open source) LibreCAD software was here first used to digitize and measure tree architecture with the SIP method. The software performed very well, providing great tools, such as precise measurement options, polyline modifications, spline through points, efficient digital image support and a convenient printing facility. Lastly, branching ratio (BR) proved to be a useful architectural trait, which may be feasibly measured with the SIP method. The two measured diameters (BT and ST) were always close to each other, and the possible inaccuracies, coming from some displacements of the measured features from the theoretical projection plane, must have been reduced while calculating the ratio.

## 4 Conclusion

The study provided both qualitative and quantitative architectural analyses of thirty post-mature temperate oaks’ forking habit; growing within a small, north-exposed forest remnant. A set of four possible qualitative models was preliminarily assumed, and finally extended to six models. These were based on the original Rauh’s HO model, differing in the location of branches that contributed to the horizontal crown extent, and including forking of the main axis. Two of the models were most clearly represented by the studied oaks. It was found that the trees tended to either keep branches at varying heights, with no forks, or to iterate forking, with no major (non-fork) branches below the first fork. The two architectural patterns resemble other HO models, such as the Attim’s model (with diffused branching) for non-forked trees, and the Leeuwenberg’s model (with equivalent branching) in case of the forked trees. The quantitative analysis confirmed the applicability of the branch to parent stem diameter ratio to define a fork; however, a 13% disagreement was found between the qualitative and quantitative fork classifications. Branching ratio was positively correlated with both tree diameter and height of a branch above the ground, which is consistent with the previous study of Colin et al. (2012), based on much younger trees. It is concluded, that most probably the tree-level factors and phenomena, such as water supplies and posture control, played the key role in the studied oaks forking habit.

---

**Fig. S1.**
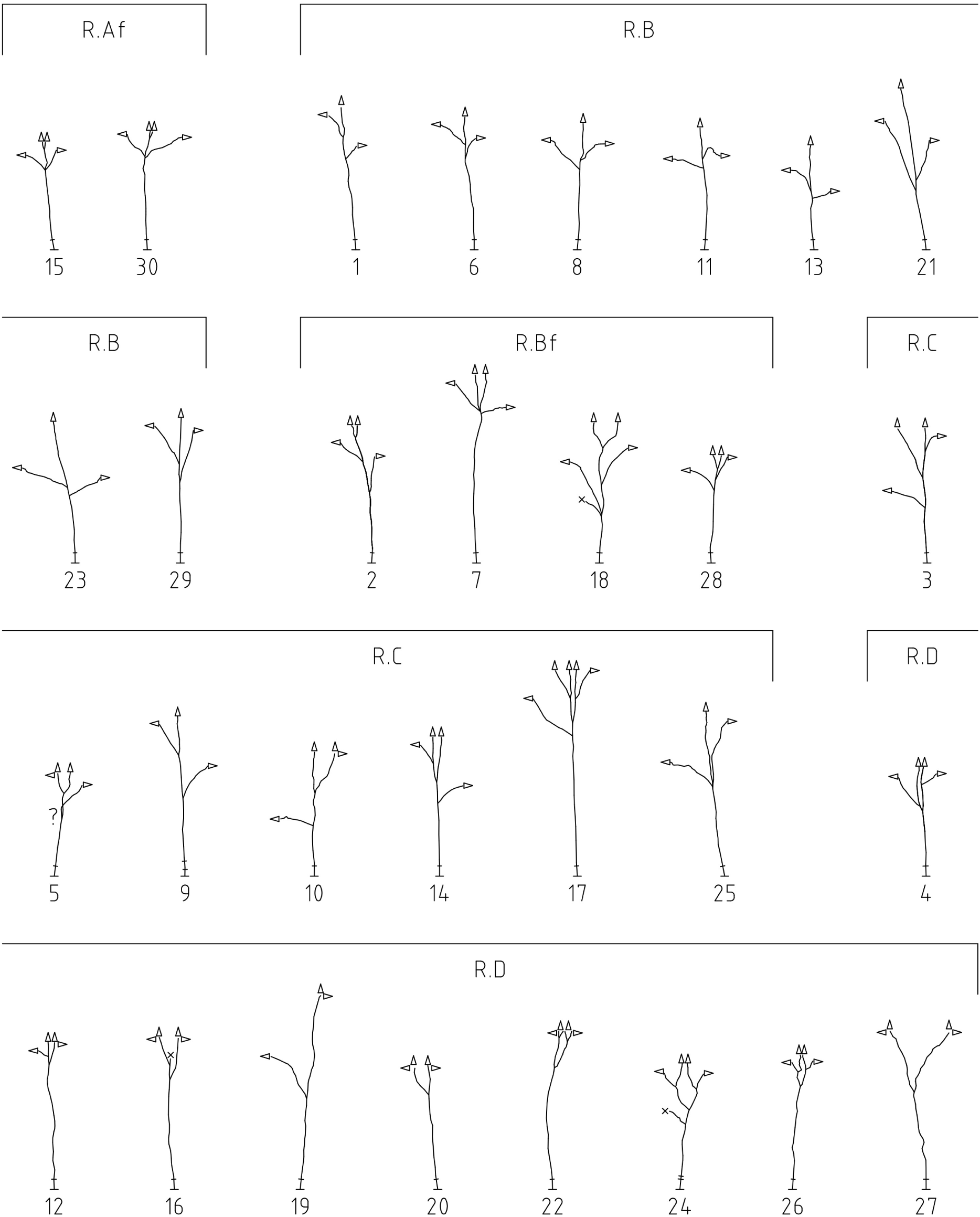
The thirty trees’ main construction (main axes and main branches), assigned to the presented qualitative models (R.A-D); the pointers indicate whether an axis contributes to the vertical, horizontal or both directions of crown extent; the short lines, just above the stem bases, represent stem diameters

**Fig. S2.**
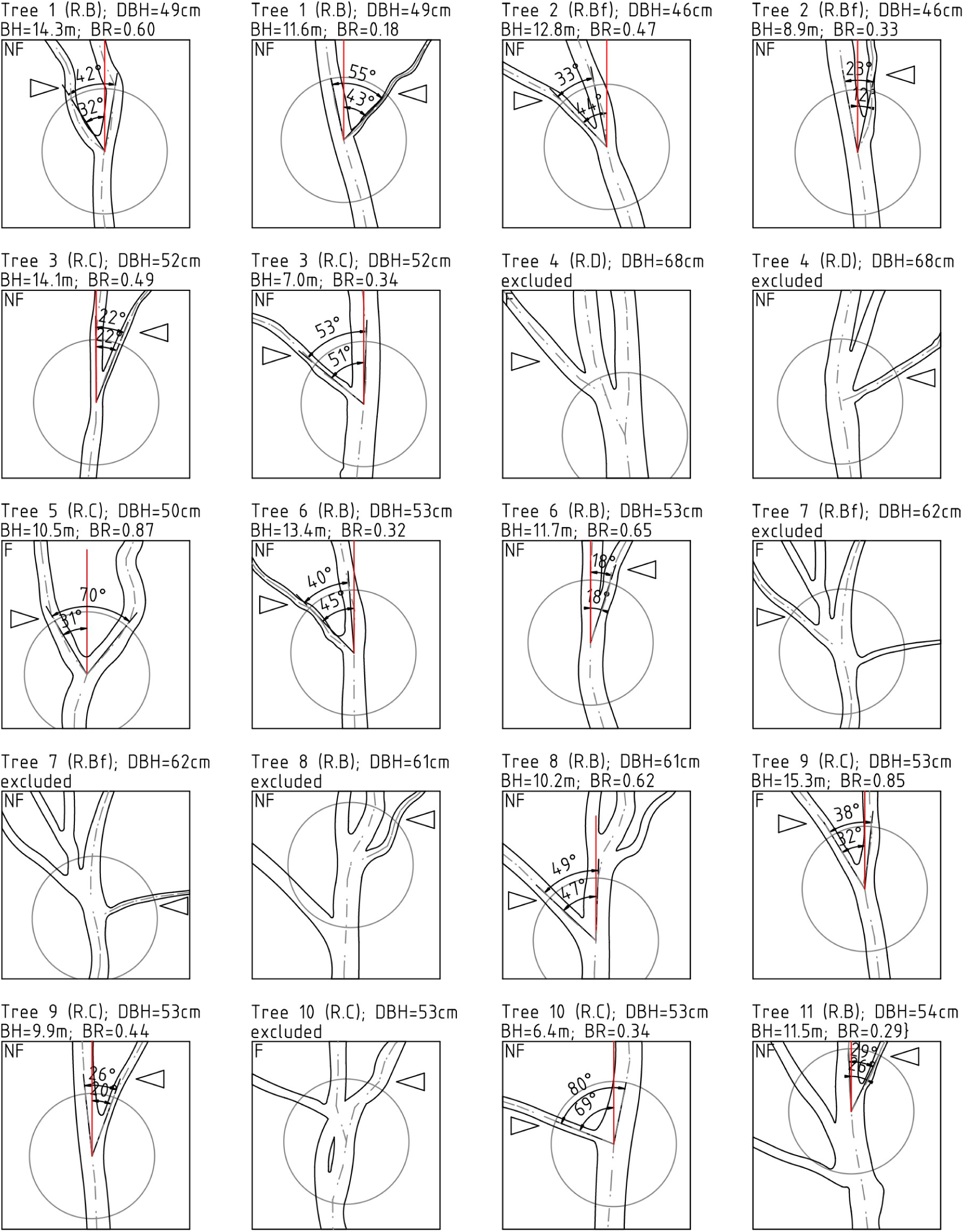
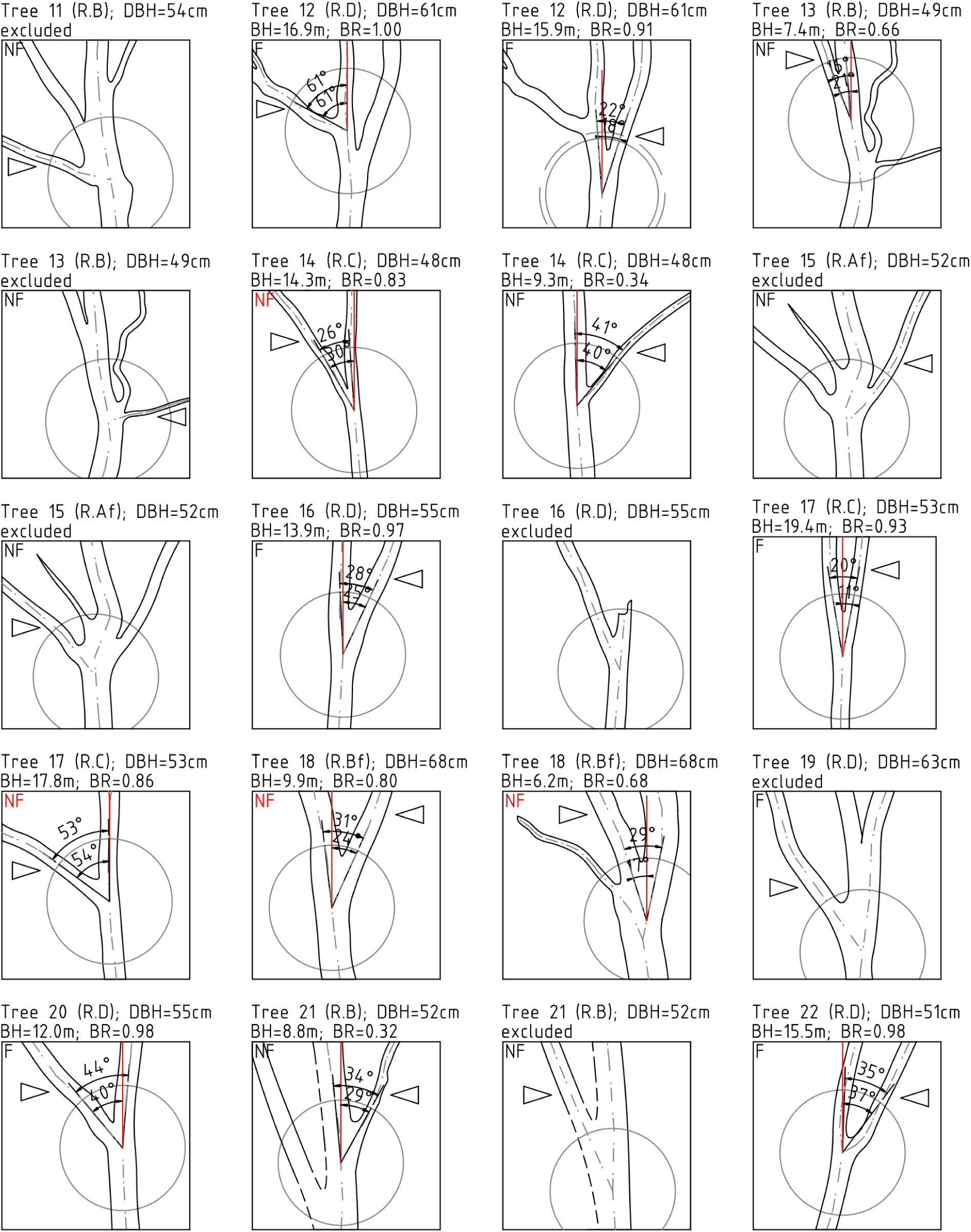
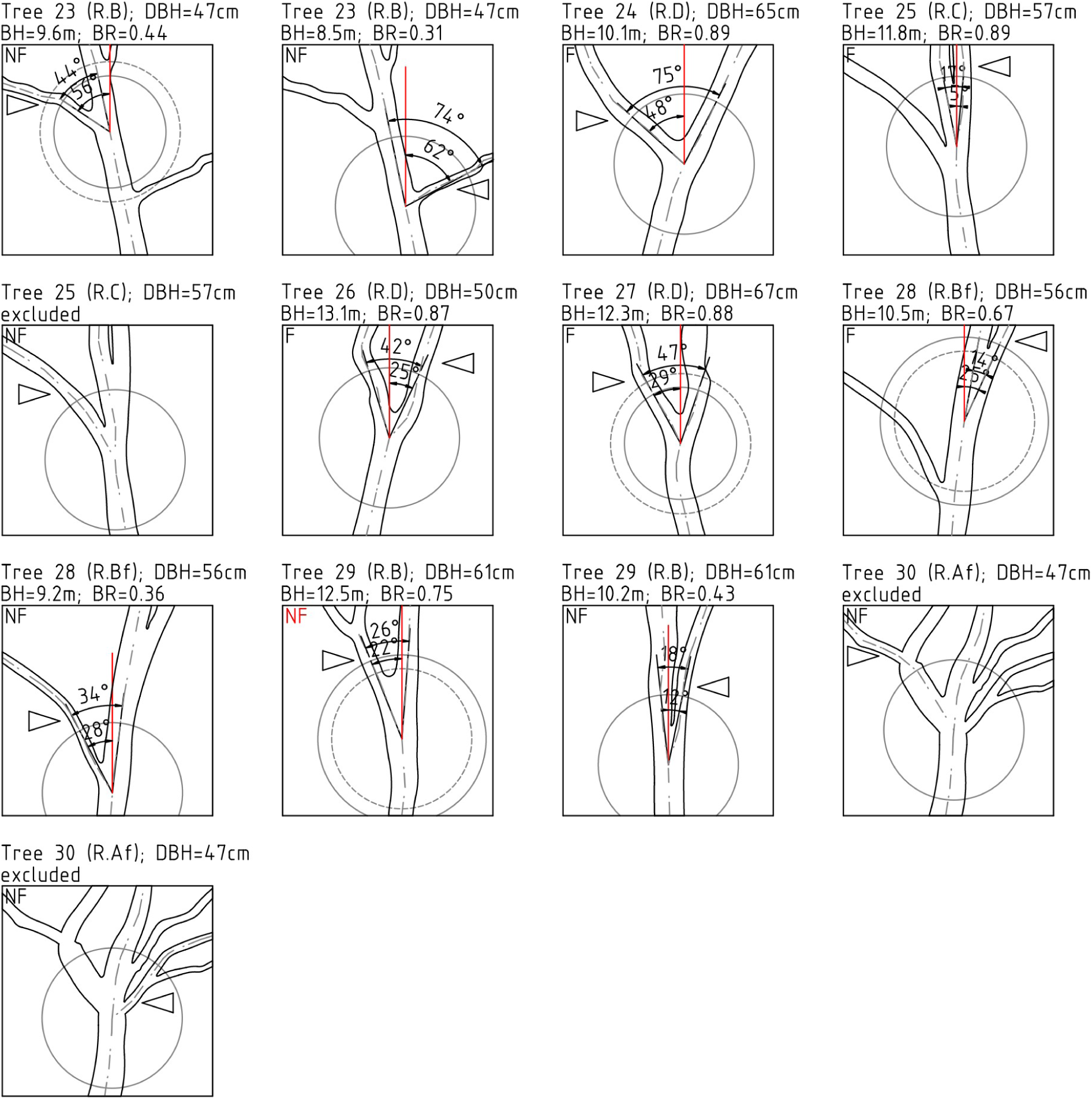
The fifty three branches (marked with triangles) analysed with the SIP method; the descriptions include: tree-level data (top lines): tree number (1 to 30), qualitative model (R.A-D), diameter at breast height (DBH); branch-level data (bottom lines): branch height (BH) and branching ratio (BR) or “excluded” if the branch was excluded from the quantitative analysis (mostly when there was another branch or branches at the distance of 1 m from the target branch insertion point); the letters in the top-left corners of the bounding boxes (3 × 3 m each) indicate qualitative branch assessments: fork (F) or non-fork (NF): in red when there was an ambiguity between quantitative and qualitative results (in five cases)

